# Validating a 14-drug microtitre plate containing bedaquiline and delamanid for large-scale research susceptibility testing of *Mycobacterium tuberculosis*

**DOI:** 10.1101/244731

**Authors:** Paola MV Rancoita, Federica Cugnata, Ana Luíza Gibertoni Cruz, Emanuele Borroni, Sarah J Hoosdally, Timothy M Walker, Clara Grazian, Timothy J Davies, Timothy EA Peto, Derrick W Crook, Philip W Fowler, Daniela M Cirillo, the CRyPTIC Consortium

## Abstract

UKMYC5 is a 96-well microtitre plate designed by the *Comprehensive Resistance Prediction for Tuberculosis: an International Consortium* (CRyPTIC) to enable the measurement of minimum inhibitory concentrations (MICs) of 14 different anti-TB compounds for >30,000 clinical tuberculosis isolates. Unlike the MYCOTB plate, on which UKMYC5 is based, the plate included two new (bedaquiline and delamanid) and two repurposed (clofazimine and linezolid) compounds. UKMYC5 plates were tested by seven laboratories on four continents using a panel of 19 external quality assessment (EQA) strains, including H37Rv. To assess the optimal combination of reading method and incubation time, MICs were measured from each plate by two readers using three methods (mirrored-box, microscope and Vizion™ Digital viewing system) after 7, 10, 14 and 21 days incubation. In addition, all EQA strains were whole-genome sequenced and phenotypically characterized by 7H10/7H11 agar proportion method (APM) and MGIT960. We conclude that the UKMYC5 plate is optimally read using the Vizion™ system after 14 days incubation, achieving an inter-reader agreement of 97.9% and intra- and inter-laboratory reproducibilities of 95.6% and 93.1%, respectively. The mirrored-box had similar reproducibilities. Strains classified as resistant by APM, MGIT960 or the presence of mutations known to confer resistance consistently record elevated MICs compared with those strains classified as susceptible. Finally, the UKMYC5 plate records intermediate MICs for one strain which the APM measured MICs close the applied critical concentration, providing early evidence that the UKMYC5 plate can quantitatively measure the magnitude of resistance to anti-TB compounds due to specific genetic variation.

## INTRODUCTION

The proportion of tuberculosis (TB) cases that are multi-drug resistant (MDR) is increasing worldwide. Although set against a background of a falling global incidence of TB, the net effect is that the number of MDR-TB cases continues to grow (1). Improving the treatment success rate for MDR-TB requires each patient to receive an individual antimicrobial regimen tailored to maximize efficacy whilst minimizing toxicity; this necessitates being able to measure minimum inhibitory concentrations (MIC) to direct both the choice of drug and dose. Universal access to prompt and comprehensive drug susceptibility testing (DST) is therefore a key component of the WHO’s *End TB Strategy* (2, 3). Although molecular approaches have the potential to deliver universal DST methods, they require further development work and any resulting solutions are likely to be expensive.

Liquid and solid media assays that measure MICs for TB exist (4–9), but are time consuming and, often, costly and so far they are not yet endorsed by any international regulatory authorities, other health international organizations such as the WHO. Microtitre plates offer a way of testing in parallel the effectiveness of a large number of drugs at a range of concentrations on small aliquots taken from a single clinical isolate. Broth microdilution methods, including several using colorimetric indicators, have previously been developed that assess the MICs for a panel of compounds using a single microtitre plate (5, 10, 11). A dry-format, 96-well microtitre plate assay (the Sensititre™ MYCOTB plate; Thermo Fisher Scientific Inc., USA) containing 12 drugs has been commercially available since 2010 and early validation studies have returned promising results (12–17). At present however, neither clinical breakpoints nor epidemiological cutoffs have been defined for any microtitre plate. Nor have any plate-based assays included both new and re-purposed drugs that will be key to the successful treatment of individual MDR-TB cases in the future.

We designed the UKMYC5 plate, which is a variant of the MYCOTB plate, to enable the Comprehensive Resistance Prediction for Tuberculosis: an International Consortium (CRyPTIC) to measure the minimum inhibitory concentrations (MICs) of 14 different anti-TB compounds (Fig. S1, Table S1) for a large number (>30,000) of clinical TB isolates that are being collected globally by participating laboratories between 2017 and 2020. Each isolate will also have its genome sequenced and the ultimate goals of the CRyPTIC project are to (i) uncover all variation in genes known to be involved in resistance and classify magnitude of their effects on specific anti-TB compounds and (ii) identify new genes that are associated with resistance. The MICs need, therefore, to be quantitatively reproducible and accurate.

In this paper, we establish the reproducibility and accuracy of the dry-form 96-well microtitre plate UKMYC5 plate for use in large-scale measurement of minimum inhibitory concentrations by the CRyPTIC tuberculosis research project. We shall assess, therefore, both the reproducibility of MIC measurements using this microtiter plate and its accuracy by comparing it to a range of established DST methods. Since the UKMYC5 plate contains 14 drugs, including in contrast to the MYCOTB plate, two new compounds (delamanid and bedaquiline) and two re-purposed drugs (clofazimine and linezolid), it could potentially, in future, form the basis of a new DST protocol for tailoring regimens to treat individual cases of MDR-TB.

## RESULTS

### Plate design and success criteria

The UKMYC5 plate was designed by the CRyPTIC consortium. The main differences compared to the commercially available MYCOTB 96-well microtitre plate are that bedaquiline, delamanid, clofazimine, and levofloxacin are included and ofloxacin, streptomycin and cycloserine were removed. The wells for each drug are arranged in a doubling dilution series. Since the plate comprises 12 columns of 8 wells and 14 drugs were included, most drugs had 6 (64x concentration range) or 7 wells, with a few having 5 or 8 (Fig. S1, Table S1).

To validate the UKMYC5 plate for use in large-scale research susceptibility testing, it is necessary to demonstrate that the plate is both reproducible and accurate. Unfortunately, no standards for Mycobacterial antimicrobial susceptibility testing (AST) exist yet. We shall therefore borrow, where appropriate, the definitions established for aerobic bacteria. The conclusions we draw about reproducibility and accuracy should therefore be treated as tentative as they will need revising once a standard for Mycobacterial AST is established.

The concentration ranges for each drug on the UKMYC5 plate are defined by the plate design and are therefore fixed. This creates a potential difficulty for assessing reproducibility and accuracy since when a strain does not grow in all the wells of a single drug, or instead grows in all the wells, the plate does not uniquely determine the MIC, it merely provides an upper and lower bound, respectively: this is termed off-scale growth. Since MICs are conventionally defined as being in agreement if they lie within a doubling dilution, then reproducibility and accuracy are only exactly defined for on-scale growth; including off-scale growth could alter the levels of reproducibility and accuracy.

### Study design

In brief, each of the seven participating laboratories (Fig. 1A) received nineteen external quality assessment (EQA) strains which were selected to ensure there was at least one strain that was resistant to each of the 14 drugs on the UKMYC5 plate (except linezolid, Table S16). Samples from each vial were sub-cultured either 2 or 10 times (Fig. 1B) and then inoculated onto a UKMYC5 plate (Fig. 1C). Each plate was incubated for 21 days, and two laboratory scientists independently determined MICs for all 14 anti-TB drugs at four different timepoints (days 7, 10, 14 and 21) using Vizion™, mirrored box and inverted-light microscope (Fig. 1D). In addition, a photograph was taken at each timepoint using the Vizion instrument, and this was retrospectively analyzed using some bespoke plate-reading software, AMyGDA (Fig. S2) (18). For more information on the above, please see the Materials and Methods.

**FIG 1.**
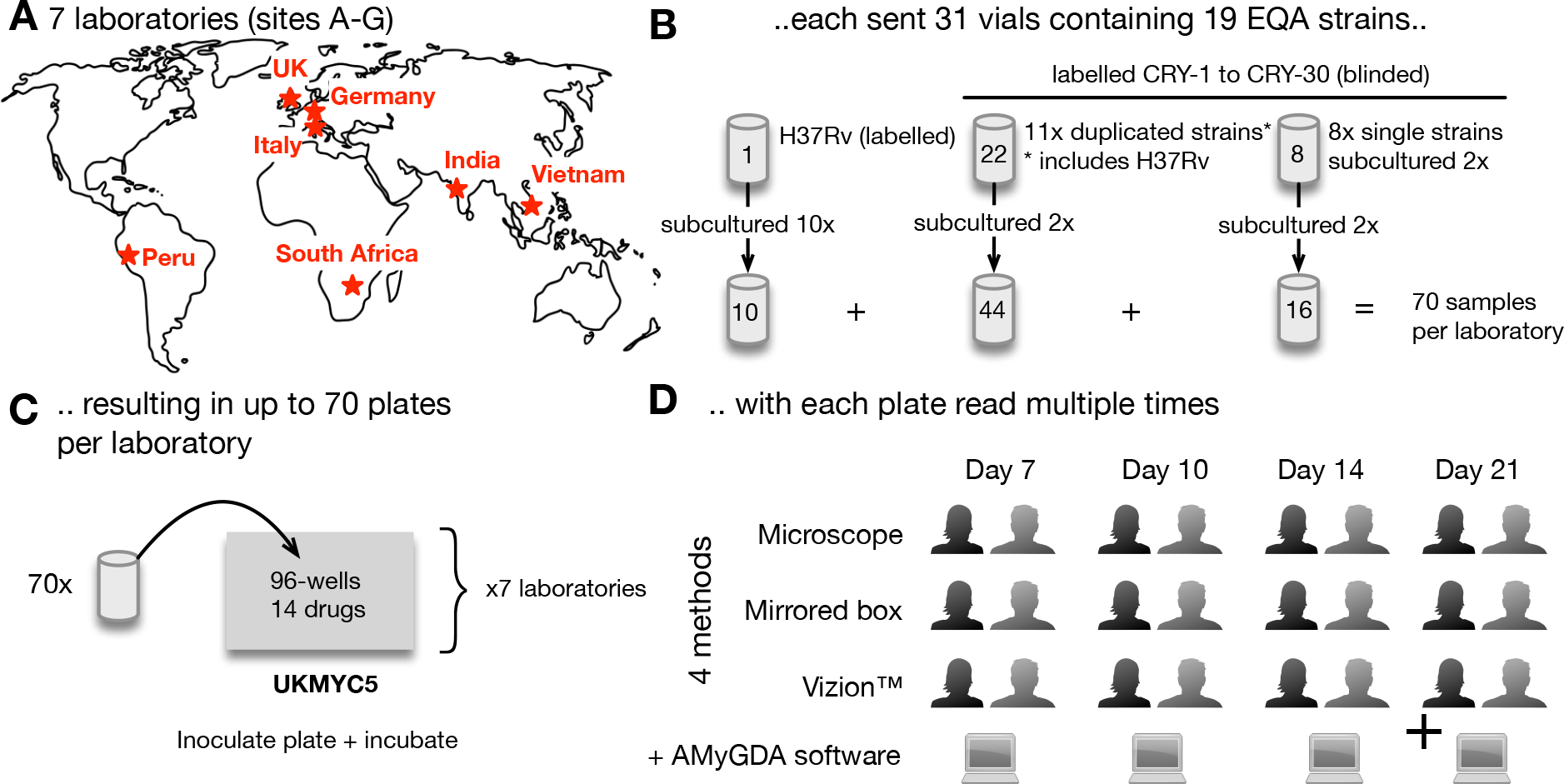
The study design for validating the UKMYC5 plate. (**A**) Seven laboratories (**B**) were each sent 31 vials containing 19 different genotypically characterized strains. H37Rv ATCC27294 was sub-cultured and tested ten times, while other strains were sub-cultured in duplicate. (**C**) Each strain was inoculated onto a UKMYC5 96-well plate starting from independent bacterial suspensions. Due to the large numbers of strains, the culturing and inoculation was performed over a period of weeks in each laboratory. (**D**) The minimum inhibitory concentrations for each drug were independently read at 7, 10, 14 & 21 days post-inoculation by two laboratory scientists using three methods as long as positive controls wells showed acceptable and visible growth. Each plate was also photographed and the image analyzed using the AMyGDA software (Fig. S2).

### The proportion of readable plates

The proportion of readable results increased with elapsed time since inoculation. As noted elsewhere (see Materials and Methods), the results for Site F were anomalous and this laboratory was excluded from the study. For the remaining six sites, the proportion of readable results was 57.8-66.1% (depending on reading method) after 7 days of incubation, which then increased to 85.7-93.0% after 14 days, reaching ≥ 95.9% after 21 days (Fig. 2A, Table S2). If a plate was not readable, this was usually because there was insufficient growth of *M. tuberculosis* in both positive control wells. As expected, the proportion of measured MICs that were off-scale in general fell as the incubation time increased (Table S4), however, even after 21 days there remained large differences between drugs, with linezolid having the fewest off-scale readings (0.4-0.8%, depending on the reading method) and rifabutin having the most (81.4-83.2%, depending on the reading method). The exact pattern for each drug depends on the plate design and how many EQA strains are resistant to its action. A logistic mixed-effects model demonstrated that significantly fewer plates were readable at day 7 and 10 than at day 14, and that significantly more plates were readable at day 21 compared with day 14 (p<0.001 for all comparisons, Table S5). The proportion of results readable by inverted-light microscope or mirrored box was significantly lower than for Vizion™ (p<0.001 in both instances, Fig. 2A, Table S5).

**FIG 2.**
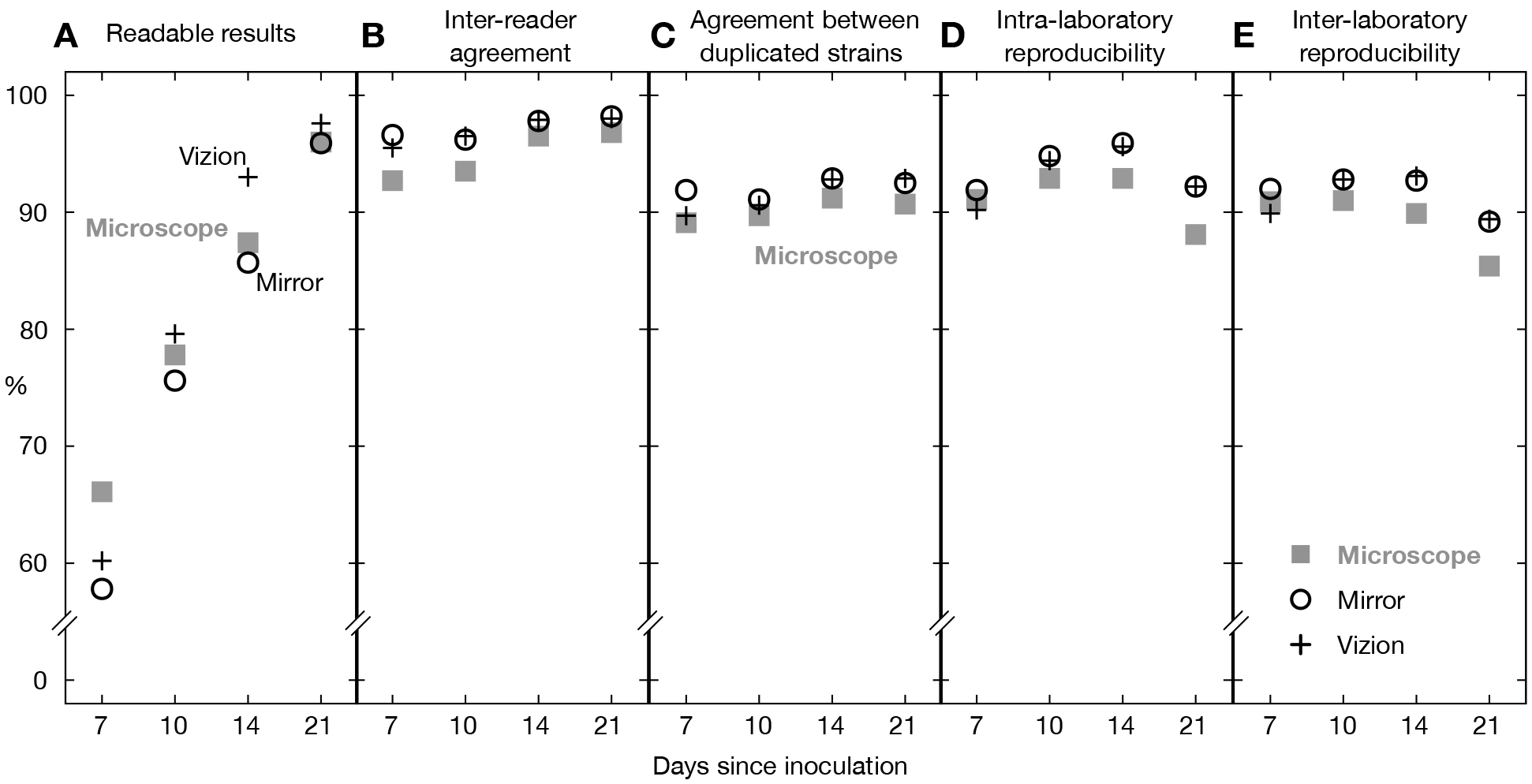
For each reading method, (**A**) the percentage of readable results, (**B**) the interreader agreement, (**C**) the agreement between two plates inoculated with the different sub-cultures of the same strain in a single laboratory and the (**D**) intra- and (**E**) interlaboratory reproducibility (Table S2). These data include both on- and off-scale MICs. Site F is excluded from this analysis.

### Agreement between readers and duplicated strains

As stated in the Materials and Methods, we define two measurements as agreeing if they are within a doubling dilution of one another. At least 92.7% of all MICs read from the same UKMYC5 plate by two scientists in the same laboratory were within one doubling dilution of one another, regardless of reading day or method (Fig. 2B, Table S2). The inter-reader agreement increased with incubation time (Fig. 2B). For Vizion™ and mirrored box, this was ≥ 95.5% across all reading-days, while for the inverted-light microscope it increased from 92.7% at day 7 to 96.8% at day 21 (Fig. 2B). The corresponding statistical model showed there was significantly lower agreement between readers when they used the inverted-light microscope compared to the Vizion™, or when they read the plates at day 7 compared to day 14 (p<0.001 for both comparisons, Table S5). The levels of inter-reader agreement were similar if we only considered the strain/drug combinations where the mode MIC was on-scale (Table S3).

As expected, the agreement between MICs measured from two UKMYC5 plates in the same laboratory, each of which having been inoculated from different cultures of the same strain taken from the same vial (i.e. duplicated strains), is lower, with ≥ 89.7% of MICs in agreement, regardless of reading-day and method (Fig. 2C, Table S2). The maximum values were observed using the Vizion instrument after 14 and 21 days incubation (92.8 and 92.9%, respectively) and with the mirrored-box after 14 days incubation (92.9%). The corresponding logistic mixed-effects model showed that the agreement of the results between duplicated strains was significantly lower at day 7 with respect to day 14 (p<0.001) and it was also lower when the results were obtained from microscope with respect to Vizion (p=0.010, Table S5).

### Reproducibility

The intra-laboratory reproducibility (the proportion of MICs within a doubling dilution of the mode, including both on- and off-scale readings) was ≥ 88.1% (Fig. 2D, Table S2) regardless of reading-day and method. The maximum values were observed after 14 days of incubation, when the mirrored-box and Vizion intra-laboratory reproducibilities were 95.9% and 95.6%, respectively. If we remove, for each drug, the strains where the modal MIC is off-scale, then the intra-laboratory reproducibility increased slightly for all reading day and methods, with maximum values of 96.5% and 96.9% for mirrored-box and Vizion, respectively, at day 14 (Table S3).

The inter-laboratory reproducibility was slightly lower, as one would expect, and was ≥ 85.4% regardless of reading-day and method combinations, peaking again after 14 days at 93.1% and 92.7% when using the Vizion system and the mirrored box, respectively (Fig. 2E, Table S2). Again removing, for each drug, strains where the MIC mode was off-scale, led to a slight increase of the inter-laboratory reproducibility at days 14 and 21 for all methods, with the highest values achieved at day 14 by Vizion and mirrored box (94.9% and 94.7%, respectively, Table S3). The logistic mixed-effects models (considering both on- and off-scale measurements) confirmed that intra- and inter-laboratory reproducibility for the inverted-light microscope was overall significantly lower with respect to Vizion™ (p<0.001 for all, Table S6). Moreover, both types of reproducibilities were lower at days 7 and 21 with respect to day 14 (p<0.001 for all, Table S6).

### Selection of reading method

Reproducibility and agreement is maximised when using either the Vizion™ system or the mirrored-box. The corresponding logistic mixed-effects models could not distinguish the performances between these two methods. However, the percentages of readable results were higher for Vizion than for mirrored-box and this was confirmed by the corresponding logistic mixed-effect model. Thus, we selected for all subsequent analyses the Sensititre™ Vizion™ Digital MIC Viewing System. Moreover, this method has the advantage that it records an image of the growth on the UKMYC5 plate which is not only a useful audit trail but also permits the use of automated plate reading software.

### Individual drug performance and selection of reading-day

To determine if the individual drugs could be read after 14 days incubation, or whether there was large variation between compounds, we analysed the individual MICs measured using Vizion™ for each drug and all reading-days (Fig. 3 & S3, Table S7). Para-aminosalicylic acid (PAS) showed the lowest inter-reader agreement at each reading day and the lowest intra- and inter-laboratory reproducibility across all reading days except day 7 (Fig. S3). This is consistent with a previous study where PAS had the lowest reproducibility amongst the 12 drugs on the MYCOTB plate (13). After 14 days incubation, all 13 remaining anti-TB compounds had values for the inter-reader agreement and intra- and inter-laboratory reproducibilities of ≥ 97.7%, ≥ 93.4% and ≥ 87.1 %, respectively. Mirroring the whole plate trends, the inter-reader agreement after 21 days tended to be slightly higher (with the exception of clofazimine, delamanid, ethionamide and kanamycin, Fig. S3), whilst both measures of reproducibility tended to fall slightly between days 14 and 21, particularly for bedaquiline, delamanid and, to a lesser extent, ethambutol (Fig. S3). Considering, for each drug, only the strains with the MIC mode on-scale most of the above trends are maintained (Table S8), with the greatest difference observed for rifabutin and delamanid, both of which had relatively few on-scale readings since in both cases only one strain out of the 19 gave an MIC mode that was on-scale. The results of the corresponding logistic mixed-effects models considering all the measurements (Table S9) showed that, overall, intra- and interlaboratory reproducibilities were significantly lower at days 7 and 21, compared to day 14 (p=0.001 and p=0.004, respectively for day 7, and p=0.001 and p<0.001 respectively for day 21). The intra- and inter-laboratory reproducibilities were indistinguishable between days 10 and 14 (p=0.465 and p=0.784, respectively) and the inter-reader agreement was lower at day 7 with respect to day 14 (p=0.030).

**FIG 3.**
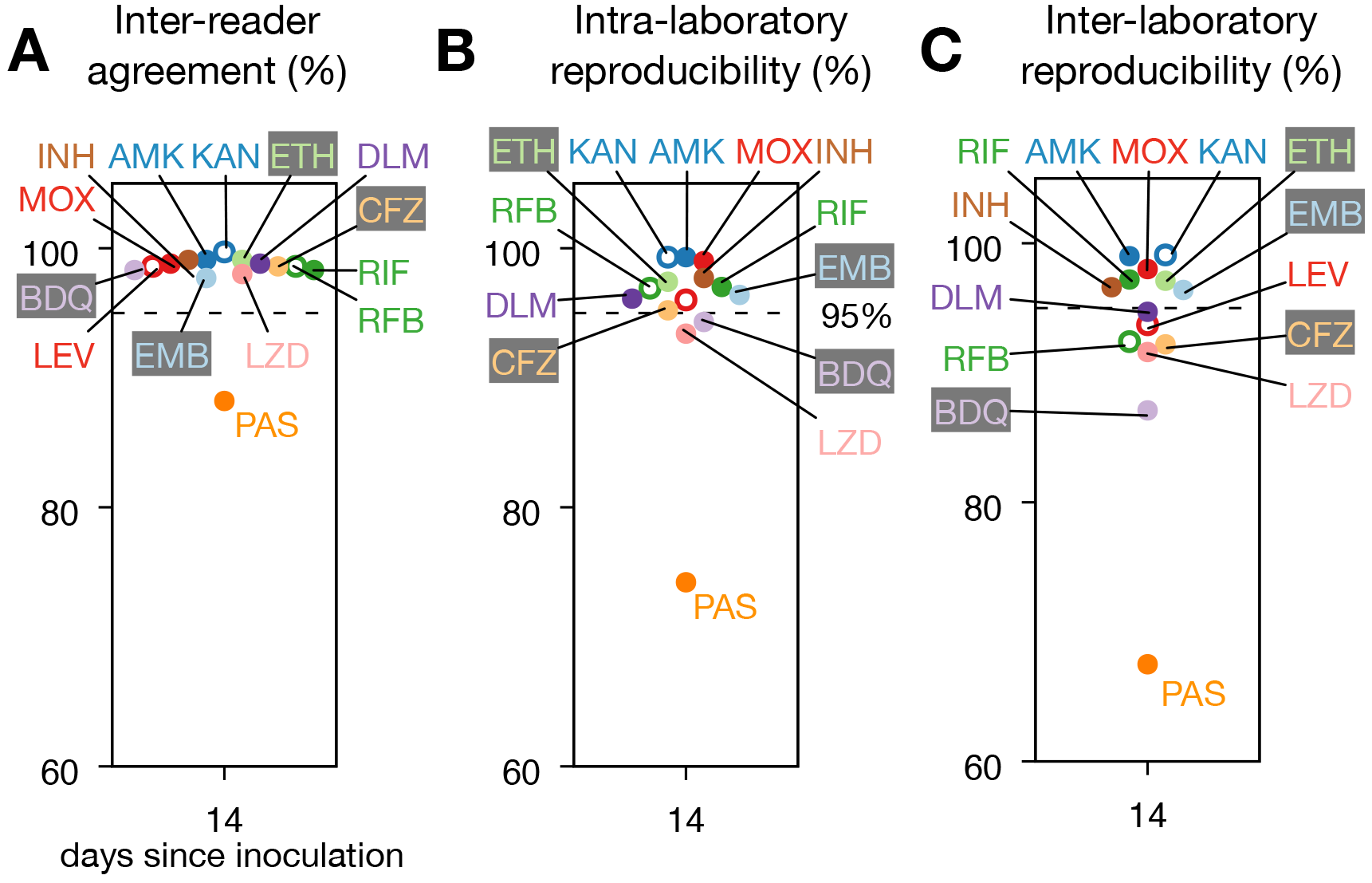
Most drugs perform well on the UKMYC5 plate, however *Para*-aminosalicylic acid (PAS) is consistently anomalous. (**A**) The inter-reader agreement for the 14 different anti-TB drugs for MICs measured after 14 days incubation using the Vizion™ system.. The (**B**) intra- and (**C**) inter-laboratory reproducibilities. A dashed line is drawn at 95%. Results for all reading days can be found in Fig S3 and Table S7.

Using the same model, reproducibility using Vizion™ was also assessed on an individual drug basis using rifampicin as a comparator selected on the basis of a known and characterised mutation in *rpoB* gene (Ser450Leu) leading to high MICs clearly distinguishable from wild-type genotypes (Table S9). Although there were statistically significant differences between many of the drugs and rifampicin, the greatest difference was seen for *para*-aminosalicylic acid where the inter-reader agreement and the intra- and inter-laboratories reproducibilities were both significantly lower than rifampicin (p<0.001). In general, although the intra- and inter-laboratory reproducibilities of the drugs were similar after 10 and 14 days (Table S9), since more results were readable at day 14 than at day 10 (Fig. 2), we elected to read the plates after 14 days of incubation.

### Automated plate reading using the with AMyDGA software

Automated plate reading software offers the ability to add a layer of quality control to plate reading, potentially increasing both reproducibility and accuracy. MICs measured from photographs of the UKMYC5 plates after 14 days incubation using the AMyGDA software (Fig. S2) (18) were then compared with MICs measured by the laboratory scientist using the Vizion™ system. As before, we considered the MICs to be in agreement if they were within a doubling dilution of one another. The agreement between the two methods starts at 87.9% at day 7 and increases with incubation time, reaching 93.8% after 21 days (Fig. S4, Table S10). At day 14, the agreement between readings by AMyGDA and Vizion™ is ≥ 90% for all drugs, except for moxifloxacin (89.3%) and para-aminosalicylic acid (73.8%). Since neither technique is an established reference method, we interpret the agreement here to measure the consistency of human- and software-based reading methods. Using a logistic mixed-effect model, we found that the agreement at days 7, 10 and 21 is not significantly different from the agreement at day 14 (p=0.143, p=0.479 and p=0.525, respectively, Table S11). We shall discuss later the potential for software like AMyGDA to further improve how microtitre plates are read.

### MIC distributions for H37Rv and agreement with endorsed methods

Having determined the optimal time and method to read the UKMYC5 plate and examined the reproducibility of this approach, we shall now assess the concordance between the UKMYC5 plate and several endorsed methods. We first compared our MICs distributions for the H37Rv ATCC27294 reference strain as measured by the Vizion™ system after 14 days of incubation with MICs measured using (i) a frozen 96-well microtitre plate, (ii) the agar proportion method (APM), and (iii) the resazurin microtitre assay (REMA). The frozen-form microtitre plate MICs were taken from existing studies, where they reported the mode of the observations (15, 16, 19). For both APM and REMA the mode of in-house measurements were taken for each drug (with the exception of ethionamide and PAS, that were not evaluated).

For the majority of drugs, there was a good level of agreement between the distribution of MICs measured using the UKMYC5 plate and the three comparator measurements (Fig. 4). The agreement between the UKMYC5 mode MICs and the mode MIC taken from the previously-published frozen-form microtitre plate study is ≥ 96.8% for evaluated drugs (i.e. bedaquiline, isoniazid, clofazimine, rifampicin, levofloxacin, moxifloxacin, amikacin, linezolid, ethambutol), with the exception of kanamycin whose agreement was 92.4% (Fig. 4, Table S12). Moreover, five of them have an agreement ≥ 99.4%. We caution, however, that since the mode MIC for four of the drugs (e.g. rifampicin, rifabutin, delamanid and clofazimine) is off-scale the ‘true’ MIC could be more than one doubling dilution from the mode of the other methods, which would reduce the level of agreement. We note that after 21 days of incubation the agreement between these methods had decreased to 65.2% and 83.0% for kanamycin and ethambutol, respectively (Table S12), providing further evidence for preferring 14 days of incubation over 21 days.

**FIG 4.**
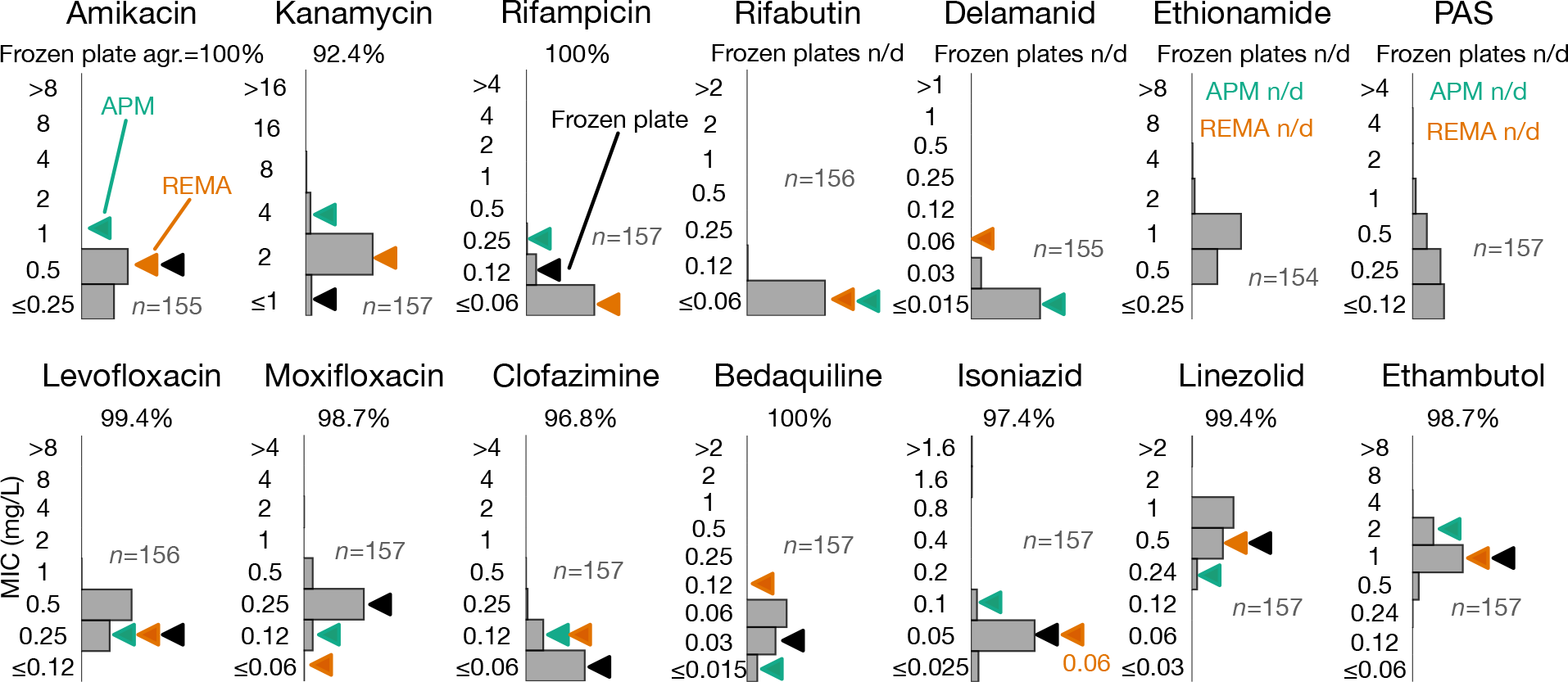
Minimum Inhibitory Concentration (MIC) distributions for H37Rv for the 14 drugs on the UKMYC5 plate, as measured at day 14 by Vizion™ are plotted as bar charts in grey. Each drug is annotated with the number of total measurements, *n*. These distributions are compared to MICs measured using three different methods: first, the published mode of the MIC for each drug (where available) measured using a frozen microtitre plate is indicated by a black filled triangle (15, 16). The agreement with the MICs measured by UKMYC5 is given for each drug. Next, the mode MICs measured by the Agar Proportion Method (APM, teal triangles) and the resazurin microtitre assay (REMA, orange triangles) are plotted. Drugs are labelled using the abbreviations defined in Table S1.

### Comparing MIC distributions of all 19 EQA strains with APM and MGIT

Since no critical concentrations exist for UKMYC5 plate, all 19 EQA strains (including H37Rv) were then categorized as resistant or susceptible using the same CC as the comparator method (as it was done in 13, 14): (i) the 7H10/711 agar proportion method (APM) and (ii) the mycobacterial growth indicator tube (MGIT, Table S22). Where possible, critical concentrations for APM were taken from recent guidance (20). The MIC distributions measured using the UKMYC5 plate were then plotted separately for the so-defined phenotypically resistant and susceptible strains for each drug (Fig. 5). For most of the drugs, resistant and susceptible strains have clearly different MIC distributions on the UKMYC5 plate (except for linezolid, which has no resistant strains, and clofazimine, while *para*-aminosalicylic acid and ethionaminde were not tested). We did not observe the lack of accuracy for ethambutol observed in previous studies using the MYCOTB plate (17, 21). Since APM and MGIT generally classified the same strains as resistant, similar distributions are observed when MGIT is used to classify the strains (Fig. 5B). The small differences between APM and MGIT arise from strains that are classified resistant by one method and susceptible by the other. For example, MGIT classified an additional strain as resistant to rifampcin which, according to APM had an MIC one doubling dilution below the critical concentration.

**FIG 5.**
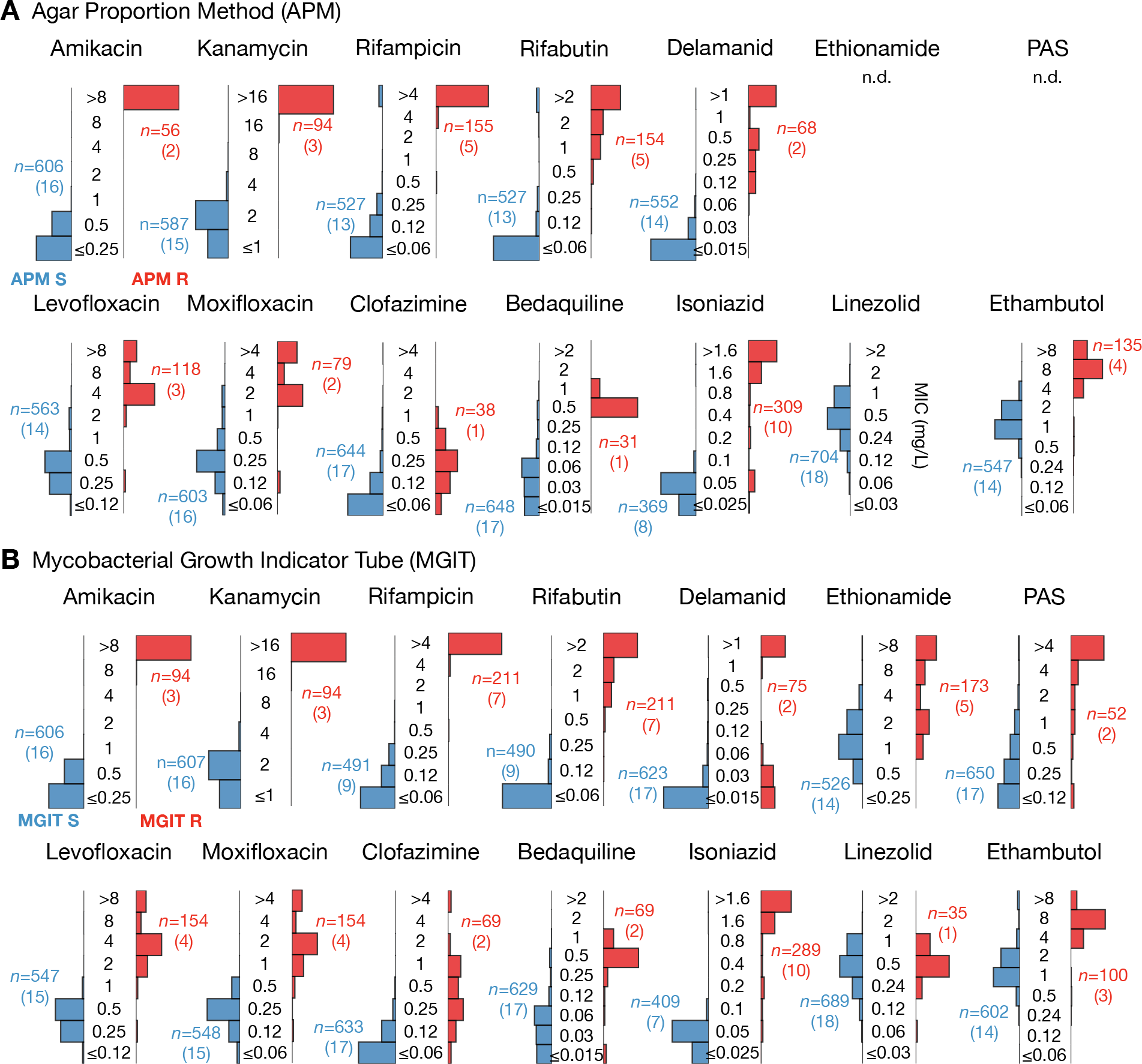
Strains identified as resistant by (**A**) the Agar Proportion Method (APM) or (**B**) the Mycobacterial Growth Indicator Tube (MGIT) consistently record elevated MICs on the UKMYC5 plate using the Vizion™ instrument after 14 days incubation. The 19 EQA strains are categorised as either Susceptible or Resistance using either APM or MGIT results. Then the distributions of MICs (mg/L) measured by all laboratories in this study (excl. Site F) for the susceptible and resistance strains are plotted in blue and red, respectively. The critical concentrations used for APM are reported in Table S13. The numbers of UKMYC5 measurements, *n*, in each bar chart is given, and in parentheses, the number of EQA strains.

To enable quantitative comparisons to be made between the UKMYC plate and both the APM and MGIT, the mode of each UKMYC5 MIC distribution was calculated by drug for each strain and then compared with the categorical results (susceptible or resistant) obtained from MGIT and APM for that strain (13, 22, 23). The categorical and conditional agreements between UKMYC5 MIC mode and those obtained from MGIT and the APM were thereby computed (Table S13). Since no critical concentrations exist for the UKMYC5 plate, we assumed shared breakpoints with each comparator method (APM or MGIT) to infer whether each strain was ‘susceptible’ or ‘resistant’ (13, 14). Discrepancies are listed in Table S14 - in the majority of cases the UKMYC5 mode was within one or two doubling dilutions of the assumed critical concentration. This could be due to the fact that the critical concentrations (CC) applied are not defined for the UKMYC5 plate. With formally defined CCs these strains may have been classified correctly.

### Comparison between genotype and UKMYC5 results

Since the genomes of all 19 EQA strains, one of which is H37Rv, are known, one can carry out a comparison between strains harbouring variations in certain genes known to confer resistance to a specific compound and those with either no variation or variation with an unknown effect or known to have no effect. For all drugs, in general, strains with genetic mutations accepted to confer phenotypic resistance to specific drugs were associated with elevated UKMYC5 MICs compared to strains with no such mutations (Fig. 6A and Table S15) (24).

**FIG 6.**
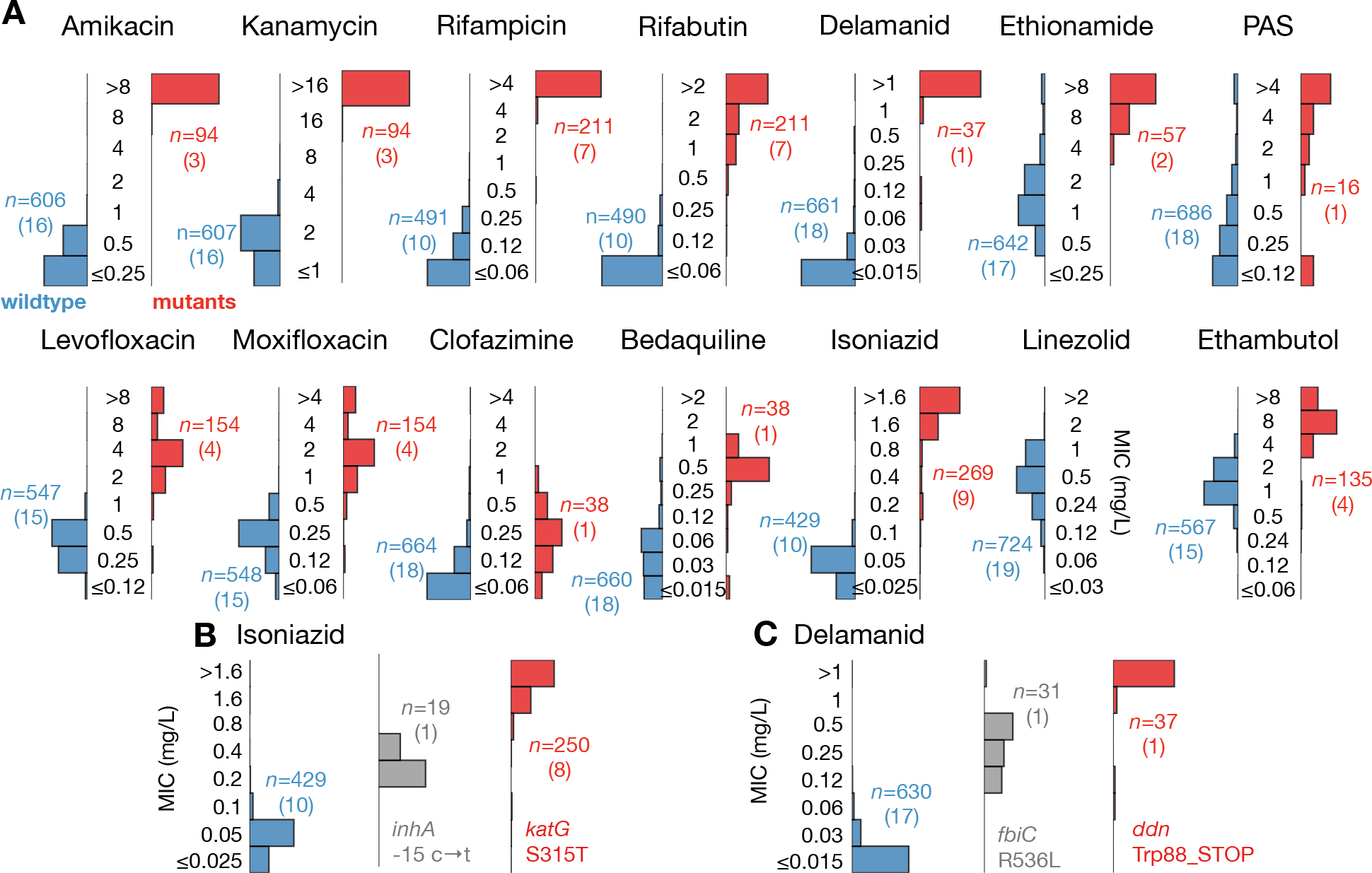
The UKMYC5 plate not only registers higher MICs for strains containing known resistance-conferring genetic variation but also reports intermediate MICs for two strains containing mutations that confer intermediate resistance to delamanid and isoniazid. (**A**) The UKMYC5 MIC distributions for each drug, split by whether the strain contains (red) or not (blue) mutations that confer resistance. Drugs that share the same resistance genes are paired. The list of genetic variants used to make this classification can be found in Table S15. The numbers of UKMYC5 measurements, *n*, in each bar chart is given, and in parentheses, the number of EQA strains. (**B**) Intermediate delamanid MICs are consistently recorded on the UKMYC5 plate for one strain containing the Arg563Leu mutation in the *fbiC* gene. (**C**) Intermediate isoniazid MICs are consistently recorded on the UKMYC5 plate for one strain containing a mutation in the promoter of *inhA*.

This is most evident for the second line injectable drugs (SLID, amikacin and kanamycin) and the rifamycins (rifampicin and rifabutin). For the SLID three of the 19 EQA strains contained the well-known a1401g mutation in the *rrs* gene (these strains were also identified by APM and MGIT as being resistant). For the rifamycins, 7 of the EQA strains contained a Ser450Leu mutation in the *rpoB* gene. The Ala90Val mutation in the *gyrA* gene, either on its own or in combination with Ser91Pro, was also associated with elevated MICs for fluoroquinolones (levofloxacin and moxifloxacin) on the UKMYC5 plate.

For isoniazid we can apply a more nuanced test since one strain contained a mutation (−15 c → t) in the promoter of *inhA* that is known to only confer an intermediate-level of resistance, whereas eight strains contained the common Ser315Thr mutation (either alone or in combination with other mutations) in the *katG* gene that is known to confer a high-level of isoniazid resistance. The promoter mutation was determined as resistant by MGIT and the APM measured its MIC to be 0.4 mg/L - just twice the critical concentration. Not only do the strains containing either of these mutations consistently have MICs higher than the remaining strains, but also the strain with the mutation known to confer an intermediate-level of resistance is observed to have intermediate-level MICs on the UKMYC5 plate (Fig. 6B) (24).

This *inhA* promoter mutation is known to also confer resistance to ethionamide (and is carried by one strain). A second strain contains an *ethA* mutation that is also accepted to confer resistance to this compound. The resulting MIC distributions for ethionamide-resistant and susceptible strains are notably cleaner and better separated than when MGIT is used to classify the strains. No strain contained a mutation known to confer resistance to linezolid. Despite some knowledge of their resistance mechanisms, the MIC distributions of the strains identified as resistant to clofazimine and *para*-aminosalicylic acid overlap with the strains assumed to be sensitive.

Interestingly, the EQA panel included two strains that are resistant to delamanid according to the APM. These have mutations in either the *ddn* gene or the *fbiC* (Rv1173) gene. The *ddn* mutation was associated with higher MICs than the *fbiC* mutation (Fig. 6C and Table S18) on the UKMYC5 plate. Consistent with this mutation having an intermediate effect, MGIT classified it sensitive and the MIC measured by APM was 0.25 mg/L - 8x less than the strain having the *ddn* mutation. Finally, four strains have a mutation in embB known to confer resistance to ethambutol (Met306Ile, Met306Val and Gln497Arg). The UKMYC5 MIC distributions for these strains are higher on average than the distribution for the remaining 15 strains.

## DISCUSSION

We have shown that it is optimal to read the UKMYC5 plate after 14 days incubation using the Thermo Fisher Sensititre Vizion™ Digital MIC viewing system. Although when using the mirrored-box the laboratory scientists were unable to read as many plates, it achieved indistinguishable levels of reproducibility to the Vizion and is cheaper so may be the preferred solution in low-income settings. The Vizion system has the added advantage of capturing a photograph of the *M. tuberculosis* growth on the UKMYC5 plate at the time of reading which can not only form a valuable audit trail, but also can be analyzed by software, providing an additional set of readings.

Using the Vizion™ system, 93.0% of UKMYC5 plates were readable after 14 days of incubation (Fig. 2). The agreement between two readers reading the same plate, or two plates containing the same strain taken from different sub-cultures, was 97.9% and 92.8%, respectively. The intra- and inter-laboratory reproducibilities were 95.6% and 93.1%, respectively. Removing strains that had potential off-scale MICs did not significantly alter these values. Considering all 14 anti-TB compounds separately, we found that *para*-aminosalicylic acid (PAS) performed poorly, consistent with a previous study (13), and the other 13 compounds individually had inter-reader agreements ≥ 97.7%, with intra- and inter-laboratory reproducibilities of ≥ 93.4% and ≥ 87.1 %, respectively (Fig. 3).

Since there are no established standards for antimicrobial susceptibility testing (AST) of *M. tuberculosis* there are no thresholds defining an acceptable level of reproducibility. To permit us to make some inferences, we shall cautiously apply the standards for aerobic bacteria, although we emphasise that any conclusions we make about the reproducibility of the UKMYC5 plate will be tentative, since they will need to be revisited when a Mycobacterial AST standard is established. For aerobic bacteria, a reproducibility of ≥ 95% is required for clinical use (25). Considering the plate as a whole, after 14 days incubation the intra-laboratory reproducibility meets this standard but the inter-laboratory reproducibility does not. Examining the drugs individually, we find that the MICs of para-aminosalicylic acid (PAS) are not reproducible after 14 days incubation with intra- and inter-laboratory values of 74.2% and 67.5%, respectively. Seven compounds do, however, satisfy the borrowed standard, having both intra- and inter-laboratory reproducibilities ≥ 95% (amikacin, ethambutol, ethionamide, isoniazid, kanamycin, levofloxacin, and rifampicin) and, with the exception of rifampicin, only considering on-scale MICs does not reduce either value below the threshold. Of the remaining six compounds, four (clofazimine, delamanid, moxifloxacin and rifabutin) have intra-laboratory reproducibilities ≥ 95% but inter-laboratory reproducibilities in the range 90 to 95%, linezolid has both reproducibilities in the range 90 to 95% and bedaquiline has intra- and inter-laboratory reproducibilities of 94.3% and 87.1%, respectively.

Assessing accuracy is more difficult since there are no critical concentrations for the UKMYC5 plate, or any other microtiter plate, thus we assumed shared breakpoints with the comparator method as was done in (13, 14). That said, we have demonstrated that the vast majority of UKMYC5 MICs for the H37Rv reference strain for each drug are within one doubling dilution of at least one, and often all three, MICs (Fig. 5) measured by APM, resazurin microtitre assay (REMA) and three studies that used a frozen-form microtitre plate (15, 16). If all 19 EQA strains are considered, we find that the UKMYC5 plate consistently and repeatedly measures elevated MICs for strains classified as resistant by APM or MGIT (Fig 5). Finally, if we instead classify the strains according to whether their genome contains mutations known to confer resistance to a specific compound, we find the UKMYC5 plate typically records elevated MICs for these strains, although we note that this analysis also makes clear our relative lack of knowledge of genetic resistance mechanisms for some compounds (notably clofazimine and PAS). Two strains contained mutations known to confer intermediate levels of resistance to isoniazid and delamanid and these strains recorded intermediate MICs on the UKMYC5 plate that were distinct from the larger susceptible and resistant populations. This is evidence that the UKMYC5 plate can also detect the magnitude of resistance.

Broth-based microdilution methods for TB DST have been proposed in the past that either directly (8) or indirectly (5, 10, 11) measure mycobacterial growth at a range of drug concentrations, potentially enabling the determination of MIC values. In 2011, the WHO examined whether non-commercial microdilution assays would be suitable to screen patients at risk of MDR-TB and conditionally recommended their use (26). Subsequent studies assessed their performance (12–14) and established MIC distributions for the H37Rv reference strain (15, 16), demonstrating a good correlation with endorsed methods.

This study moves the field forward: UKMYC5 is the first microtitre plate assay to incorporate both new (delamanid, bedaquiline) and repurposed drugs (linezolid, clofazimine) in a dry-well format that is convenient to transport and store. The results presented here indicate that the UKMYC5 plate is sufficiently reproducible for large-scale research susceptibility testing of *Mycobacterium tuberculosis* and, in particular, it appears able to detect genetic variants which confer intermediate levels of resistance, which is an important subsidiary goal of CRyPTIC. In future work we shall assess the ability of the AMyGDA software to add some quality control and, we hope, improve the reproducibility and accuracy of the MICs measured.

In its present form, however, the UKMYC5 plate is not suitable for clinical use. The overall reproducibility and accuracy of the plate would benefit from a redesign to remove PAS and the vacated wells used to expand the range of concentrations for drugs where measurements were most frequently reported at the extremes of the dilution range; this redesign is underway. We hope that by the time CRyPTIC project has built a large dataset of unbiased clinical samples using the UKMYC5 plate (and its successor) an antimicrobial susceptibility testing standard for Mycobacteria will have been established which will allow the formal evaluation of the suitability of broth-based microtitre plates for clinical use in diagnosing and treating tuberculosis.

The UKMYC5 plate does, however, have several limitations that need acknowledging; these include the need for a pre-culture step, entailing a delay of up to six weeks before the plate is inoculated. This currently prevents the rapid turn-around of results, however, we expect this time will be reduced significantly by further development work to define the optimal inoculum from MGIT culture. In addition, unlike several commercial tests, a microtitre plate such as UKMYC5 can only be used in a biosafety level 3 laboratory, which could affect its potential utility in clinical microbiology. Despite these shortcomings, the possibility of a single microtitre plate that could provide MICs for a range of anti-TB compounds at relatively low cost should be vigorously pursued.

The determination of MIC values for a range of drugs allows therapeutic decisions to be more nuanced since they can be guided by the degree of resistance to a drug and an understanding of the tolerability of drug doses required to overcome it. Unlike CC-based DST methods, where all errors are categorical in nature, MIC errors can be marginal, and thereby may be potentially less disruptive to treatment decisions. In addition to these advantages, the UKMYC5 plate assays 14 drugs at once, at low cost - a clear advantage over MGIT and the APM. Including both new and re-purposed drugs is a clear advantage over other microtitre plate-based quantitative DST methods.

As different countries have different settings, it would be optimal if one could modify the design of the microtitre plate to reflect what drugs are locally available, or the make-up of locally recommended regimens. It is therefore key that, whilst the ranges of MICs are standardised, the combination of drugs included on the plate can be adjusted according to need and this does not significantly increase the manufacture cost. A challenge to regulators is therefore whether the performance of the plate can be accredited for a menu of *individual* drugs from which country-specific plates can be constructed, rather than a fixed plate layout.

Since the increasing global numbers of MDR-TB cases are a major obstacle to TB control, let alone its elimination, there is growing need for a clinical assay that can provide quantitative data on the second-line, new and repurposed drugs, clinicians will have to prescribe with increasing frequency in the future. The UKMYC5 plate and its successor has the potential to guide the appropriate treatment of MDR-TB directly through implementation in clinical microbiology laboratories fulfilling all biosafety requirements for handling MTBC isolates or by providing detailed and timely surveillance of the prevalence of different strains by country. The Foundation for Innovative and New Diagnostics (FIND) have already expressed interest in pursuing endorsement by the WHO of the UKMYC5 plate. In parallel, its impact on our understanding of the effect of genomic mutations on the MICs of various drugs is likely to inform the WHO’s planned assessment of WGS-based DST in 2018 (1). Whether it be through direct measurement of MICs, or predictions of MIC from genomic data, a new era of quantitative TB DST may be almost with us.

## MATERIALS AND METHODS

### Participating laboratories

Thirty one vials containing nineteen external quality assessment (EQA) TB strains were distributed by the WHO Supranational Reference Laboratory at San Raffaele Scientific Institute (SRL), in Milan, Italy, to six other participating laboratories (Fig. 1A). These were located in Germany (Institute of Microbiology and Laboratory Medicine, IML red GmbH, Gauting), UK (Public Health England Regional Centre for Mycobacteriology, Birmingham), India (P.D. Hinduja National Hospital and Medical Research Centre and Foundation for Medical Research, Mumbai), South Africa (Centre for Tuberculosis at the National Institute for Communicable Diseases, Johannesburg), Peru (Mycobacterial Laboratory, Cayetano Heredia University, Lima) and Vietnam (Oxford University Clinical Research Unit in Vietnam, Ho Chi Minh City).

### *M. tuberculosis* strains

Each participating laboratory received up to 31 culture vials of *Mycobacterium tuberculosis* (Fig. 1B & S17). One contained the H37Rv *Mycobacterium tuberculosis* reference strain ATCC 27294, purchased from BEI Resources (Manassas - Virginia) (GenBank AL123456) (27) and was labelled as such. Eleven EQA strains were sent in duplicate (one of which was also H37Rv) by the SRL, along with another eight additional EQA unique strains, bringing the total to 31 culture vials containing nineteen distinct strains. Except for H37Rv ATCC 27294, other samples were coded CRY-1 to CRY-30, and therefore were blinded (Fig. 1B).

H37Rv ATCC 27294 was chosen as the reference strain since it is susceptible to all anti-tubercular drugs on the UKMYC5 plate. The other 18 strains were chosen from clinical isolates or proficiency testing isolates stored in the repository at SRL Milan, based on the presence of mutations known to confer resistance to specific drugs with a high-degree of confidence (Table S16) and therefore could be reasonably expected to have intermediate or high MICs values on the UKMYC5 plate.

### Preparation of replicates

Upon arrival, all strains were sub-cultured onto solid media as follows: H37Rv was subcultured in 10 different Loewenstein-Jensen tubes or 7H10 agar plates while the remaining 30 vials of the validation panel were sub-cultured in duplicate onto solid media (Loewenstein-Jensen or 7H10 agar plates). Therefore each participating laboratory tested up to 70 UKMYC5 plates (Fig. 1C, S5).

### UKMYC5 design

The UKMYC5 plate was designed by the CRyPTIC consortium and manufactured by Thermo Fisher Scientific Inc., UK. Fourteen anti-TB drugs (rifampicin, rifabutin, isoniazid, ethambutol, levofloxacin, moxifloxacin, amikacin, kanamycin, ethionamide, clofazimine, *para*-aminosalicylic acid, linezolid, delamanid and bedaquiline) were included, at 5 to 8 doubling dilutions (Fig. S1, Table S1). Janssen Pharmaceutica and Otsuka Pharmaceutical Co., Ltd provided bedaquiline and delamanid pure substances, respectively. Although pyrazinamide-only plates containing lyophilised substance in different stocks were tested, poor performance due to suboptimal broth pH conditions resulted in pyrazinamide being excluded from UKMYC5. Two batches of UKMYC5 plates were manufactured and distributed for this study.

### Inoculation protocol

Scientists from all laboratories received training in plate inoculation and reading at the SRL, Milan. The standard operating procedure involved preparing a 0.5 McFarland suspension in saline tween with glass beads (Thermo Fisher, Scientific Inc., USA) from 20-25 day-old colonies (or no later than 14 days after visible growth) grown on Löwenstein-Jensen or 7H10 agar media. Duplicates were tested on different days starting from different subcultures, thus starting from different bacterial suspensions.

Suspensions were then diluted 100-fold with the addition of 100 μL of the suspension to 10 mL of enriched 7H9 broth. Aliquots of 100 μL of standard 1.5×10^5^ CFU/mL inoculum (approximate range 5×10^4^ – 5×10^5^) were dispensed into wells by the semi-automated Sensititre™ Autoinoculator (Thermo Fisher, Scientific Inc., USA). Each plate was then sealed using a manufacturer-supplied rectangle of transparent plastic that remains in place throughout the 21 day incubation period.

### Measurement of MICs

In each laboratory, two scientists independently read each microtitre plate using three different methods (Thermo Fisher Sensititre™ Vizion™ Digital MIC viewing system, a mirrored-box and an inverted-light microscope) at 7, 10, 14 and 21 days postinoculation (Fig. 1D). A plate was considered valid when both positive control wells (containing no drug) showed acceptable and visible growth. MIC results and additional data were recorded locally onto paper and into a shared web-enabled database (Table S17). An image of each plate was captured using the Vizion™ and was stored and subsequently analysed by software, the Automated Mycobacterial Growth Detection Algorithm (AMyGDA) (18). AMyGDA analysis was performed at the University of Oxford using default settings (Fig. S2).

### Laboratory Validation

For Site F the proportion of readable results was anomalously high (≥ 94.2%), regardless of reading day (Fig. S6, Table S20), yet Site F had an anomalously low intralaboratory reproducibility, varying between 77.5%-80.2% depending on the reading day (Fig. S6, Table S20). Site F also had an anomalously low inter-laboratory reproducibility of only 72.2%-75.3% (Fig. S6, Table S18). Logistic mixed-effect models confirmed that for Site F inter-reader agreement (being within a doubling dilution) and intra- and interlaboratory reproducibilities were all significantly lower than the other laboratories (p<0.001 for all comparisons; Table S19). Data from Site F was consequently excluded from subsequent analyses. A retrospective analysis of the plate images recorded by Vizion™ revealed that readers at Site F had frequently mistaken sediment from the inoculum with bacterial growth, thereby allowing us to offer targeted training to the laboratory scientists involved. This process highlights the importance of storing images of all the plates to form an audit trail.

### Independent characterisation of panel strains

Each of the 19 strains used (Fig. 1B) were phenotypically characterised by BACTEC™ MGIT960 (BD Lifesciences, New Jersey, USA), Middlebrook 7H10/7H11 agar dilution method (Table S20) and (with the exception of ethionamide and *para*-aminosalycilic acid) resazurin microtitre assay (REMA, Table S21) for drugs for which the WHO has endorsed critical concentrations (CCs) (23). Middlebrook 7H11 agar was used for bedaquiline and delamanid. All strains were also whole genome sequenced; genomic DNA was extracted from Löwenstein-Jensen cultures using either FastPrep24 for cell lysis and ethanol precipitation or the cetyl-trimethylammonium bromide method as described elsewhere (28). Paired-end libraries of 101 bp were prepared using the Nextera XT DNA Sample Preparation kit (Illumina Inc., San Diego, CA, USA) and sequenced on Illumina HiSeq 2500 instruments (with the exception of one site which used NextSeq 500 instruments). A minimum genome coverage of 30x was required for SNP analysis. Variant calling in genes associated with resistance was performed by the PhyResSE web tool and the bioinformatics pipeline at the SRL (29) and the results are given in Tables S17 & S18. The sequences reported in this paper have been deposited in the Sequence Read Archive of the National Center for Biotechnology Information under study accession numbers SRP068011 & SRP130092.

### Statistical analysis

Both descriptive and modeling analyses were conducted. For the latter logistic mixed-effects models were constructed since the data consist of repeated measurements. A drug on a plate is defined as readable if (i) there is acceptable growth in both the positive control wells, (ii) there is no contamination in the wells for that drug and (iii) the wells of that drug were not evaporated. We define two measurements as being in agreement if the two MICs are within one doubling dilution of each other. Furthermore, the reproducibility is defined as the proportion of MICs that are within one doubling dilution of the mode. Due to the high percentage of off-scale results, results were considered within one doubling dilution if they were adjacent in the doubling dilution range even if one of them was off-scale. This constitutes a best-case approach, and therefore where appropriate calculations were repeated only using drug,strain combinations where the MIC mode was on-scale. To assess the intra-laboratory reproducibility, the mode MIC for each drug was computed for that site, pooling the results across reading methods, days, replicates and readers. To test the interlaboratory reproducibility, the mode was calculated only for each drug, pooling the results also across sites (besides reading methods, days, replicates and readers). To assess the accuracy of the plate we compared UKMYC5 MICs to the mode of the MICs obtained by other established DST methods. For both APM and REMA two independent measurements were taken for each drug (with the exception of ethionamide and PAS, that were not evaluated) and, where there was a discrepancy (i.e. the mode was not unique), the lower value was chosen. Since no critical concentrations exist for UKMYC5 plates, we assumed the same critical concentration of the comparator method (as it was done in 13, 14). Results were assessed in three ways: (i) the agreement between the UKMYC5 and an established method was calculated, (ii) the categorical agreement was defined as concordant reporting of either sensitivity or resistance as defined by the critical concentration (CC) of the comparator phenotypic test (if the MIC was lower or equal to the CC the strain was defined as susceptible, otherwise it was defined as resistant). Finally, (iii) the conditional agreement was defined as resistance by the comparator method and a MIC equal to or higher than the CC on the UKMYC5, or susceptibility by the comparator method and a MIC equal to or lower than the CC plus one doubling dilution on the UKMYC5 (14).

## ACKNOWLEDGMENTS

We acknowledge the CRyPTIC members for their hard work which made this study possible, and thank the following collaborators for their crucial roles: Marco Rossi and Giovanna Graziano from the SRL Milano for the organisation of the training sessions in Milan; Meera Gurumurthy (The Union) for the training session in Beijing; Hien Ho Van and Tien Nguyen Thanh (Oxford University Clinical Research Unit, Ho Chi Minh City) for the development and maintenance of the CRyPTIC database; Koné Kaniga (Janssen Ltd) and Yongge Liu (Otsuka Pharmaceutical) for making available bedaquiline and delamanid powder, respectively; Thermo Fisher Scientific Inc, for the manufacture of the microtitre plate assays. Finally, we thank Claudio Köser for constructive discussions.

### Members of the CRyPTIC consortium

Derrick W Crook, Timothy EA Peto, A Sarah Walker, Sarah J Hoosdally, Ana L Gibertoni Cruz, Clara Grazian, Timothy M Walker, Philip W Fowler, Daniel Wilson and David Clifton, University of Oxford; Zamin Iqbal and Martin Hunt, European Bioinformatics Institute; E Grace Smith, Priti Rathod, Lisa Jarrett^⋆^ and Daniela Matias^⋆^, Public Health England, Birmingham; Daniella M Cirillo, Emanuele Borroni^⋆^, Simone Battaglia^⋆^*, Matteo Chiacchiaretta^⋆^, Maria De Filippo and Andrea Cabibbe, Emerging Bacterial Pathogens Unit, IRCCS San Raffaele Scientific Institute, Milan; Sabira Tahseen, National Tuberculosis Control Program Pakistan, Islamabad; Nerges Mistry, Kayzad Nilgiriwala^⋆^, Vidushi Chitalia^⋆^, Nithyakalyani Ganesan^⋆^ and Akshata Papewar^⋆^, The Foundation for Medical Research, Mumbai; Camilla Rodrigues, Priti Kambli^⋆^, Utkarsha Surve^⋆^ and Rukhsar Khot^⋆^, P.D. Hinduja National Hospital and Medical Research Centre, Mumbai; Stefan Niemann, Thomas Kohl and Matthias Merker, Research Center Borstel; Harald Hoffmann, Sarah Lehmann^⋆^ and Sara Plesnik^⋆^, Institute of Microbiology & Laboratory Medicine, IML red, Gauting; Nazir Ismail, Shaheed Vally Omar, Lavania Joseph^⋆^ and Elliott Marubini^⋆^, National Institute for Communicable Diseases, Johannesburg; Guy Thwaites, Thuong Nguyen Thuy Thuong, Nhung Hoang Ngoc^⋆^ and Vijay Srinivasan^⋆^, Oxford University Clinical Research Unit, Ho Chi Minh City; David Moore, Jorge Coronel^⋆^ and Walter Solano^⋆^, London School of Hygiene and Tropical Medicine and Universidad Peruana Cayetano Heredfa, Lima; Guangxue He, Baoli Zhu, Yanlin Zhou, Aijing Ma and Pang Yu, China CDC, Beijing; Marco Schito, The Critical Path Institute, Tucson. Ian Laurenson and Pauline Claxton, Scottish Mycobacteria Reference Laboratory.

## Footnotes

* directly involved in laboratory work

## SUPPLEMENTAL MATERIAL

Supplementary material (Figures S1 to S6 and Tables S1 to S22) for this article may be found online.

